# Acyloxyacyl Hydrolase Regulates Microglia-Mediated Pelvic Pain Through Toll-Like Receptor-4

**DOI:** 10.1101/2021.02.26.433087

**Authors:** Afrida Rahman-Enyart, Ryan E. Yaggie, Wenbin Yang, Justin L. Bollinger, Deborah R. Winter, Anthony J. Schaeffer, David J. Klumpp

## Abstract

Interstitial cystitis/bladder pain syndrome (IC/BPS) is a devastating condition of chronic pelvic pain and urinary dysfunction. We have shown that mice deficient for the lipase acyloxyacyl hydrolase (AOAH) develop pelvic allodynia and exhibit symptoms and comorbidities consistent with IC/BPS, as well as gut dysbiosis. Microglia are resident immune cells of the central nervous system (CNS) that respond to changes in the gut microbiome, and studies have linked microglial activation to neuropathic pain. Additionally, microglia express toll-like receptors (TLRs), including TLR4, which are activated by microbial components. We have previously shown that AOAH-deficient mice exhibit increased gut permeability, suggesting a possible mechanism of microglial TLR4 activation via translocation of microbial products across the intestinal barrier to the brain. Here, we assessed the role of AOAH and TLR4 in microglial activation and pelvic pain. AOAH immunoreactivity co-localized with the microglial marker P2YR12 but not astrocytes, suggesting a functional role for AOAH in microglia. Pharmacologic ablation of CNS microglia with PLX5622 resulted in decreased pelvic allodynia in AOAH-deficient mice and resurgence of pelvic pain upon drug washout. Aligned with microglial activation, we observed altered cytokine abundance in *Aoah*−/− cortex that was reduced in *Aoah/Tlr4*−/− cortex. Consistent with our hypothesis of TLR4 activation by gut microbes, we observed microbiome-dependent activation of cultured BV2 microglial cells. Skeletal analyses revealed that AOAH-deficient mice have an activated microglia morphology in brain regions associated with neuropathic pain, independent of TLR4. Compared to *Aoah*−/− mice, *Aoah/Tlr4*−/− mice exhibited decreased pelvic pain and microglial cytokine expression. Together, these findings demonstrate differential roles for AOAH and TLR4 in microglial activation and pelvic pain and thus identify novel therapeutic targets for IC/BPS.

## INTRODUCTION

Interstitial cystitis/ bladder pain syndrome (IC/BPS) is a debilitating condition characterized by chronic pelvic pain and other lower urinary tract symptoms [3,28,39]. Treating IC/BPS is challenging due to diverse disease etiologies and a lack of biomarkers, making advances towards understanding IC/BPS a necessity [6]. Towards this goal, we previously conducted a genetic screen in a mouse model of pseudorabies virus-induced pelvic pain and identified a polymorphism near the gene encoding *acyloxyacyl hydrolase*, *Aoah* [61].

AOAH is a highly conserved lipase responsible for hydrolyzing the secondary fatty acyl chains of gram-negative bacterial lipopolysaccharides (LPS), resulting in LPS detoxification and attenuation of host inflammation [19,21,35,40,51]. We have previously reported that mice deficient for AOAH exhibit enhanced pelvic allodynia and are more vulnerable and responsive to induced pelvic pain models [61]. Furthermore, we have also identified AOAH deficiency as a mediator of arachidonic acid-dependent expression of corticotropin releasing factor (CRF) in the paraventricular nucleus (PVN) of the hypothalamus [2], a pertinent brain region for CRF-dependent pain modulation [31,54]. In addition, AOAH-deficient mice mimic other facets of IC/BPS, including a depressive phenotype and dysbiosis of the gut microbiome [1,2,8,12,24,38,42].

Gut microbiota and associated metabolites are key regulators of the immune response, and play an important role in the development and maturation of microglia [16,18,36]. Best known as the immune cells of the central nervous system (CNS), microglia are also fundamental for brain development and maintenance of homeostasis [33,34]. Microglial morphology is highly dynamic in response to environmental cues ranging from a ramified phenotype of resting cells in surveillance-mode to a bushy/ameboid phenotype of activated cells associated with increased cytokine release, proliferation, migration, and phagocytosis [4].

The gut microbiome has previously been shown to regulate microglial function [18,36,55], and both are well-documented in the development and maintenance of chronic pain [5,16,20,22,26]. Several mechanisms of how gut microbes may interact with microglia and regulate chronic pain have been proposed, including the leakage of bacterial products across an impaired gut barrier to the brain [16]. Previous studies have shown that bacterial LPS can directly bind to and activate microglial cells via toll-like receptor-4 (TLR4), a member of the pattern recognition receptor family [13,32,46]. Since AOAH-deficient mice exhibit compromised gut epithelia and gut dysbiosis [42], we hypothesized that AOAH deficiency would result in microglial activation, dependent on TLR4. We observed that AOAH-deficient mice exhibited activated microglia in brain regions important for pain modulation, and activation was dependent on the presence of TLR4. Furthermore, we observed that microglia regulated pelvic pain in AOAH-deficient mice, as microglial ablation partially alleviated the pelvic pain phenotype. These findings suggest that microglial activation can regulate pelvic pain and provide insights for future development of therapeutics for pelvic pain disorders such as IC/BPS.

## MATERIALS AND METHODS

### Animals

Eight to ten-week-old female wild type (WT) C57BL/6 mice were purchased from The Jackson Laboratory. *Aoah*−/− mice (B6.129S6-Aoahtm1Rsm/J) were a generous gift from Dr. Robert Munford of NIAID and maintained as previously described [2]. *Aoah*−/−/*Tlr4*−/− mice were generated by intercrossing *Aoah*−/− mice with *Tlr4*−/− mice (The Jackson Laboratory, Bar Harbor, ME) followed by intercrossing of F1 to get F2 and genotyping to ensure double knockout.

### Immunohistochemistry

Coronal brain sections for immunohistochemistry were prepared from WT, *Aoah*−/−, and *Aoah*−/−/*Tlr4*−/− mice. The mice were first anesthetized with isoflurane and then transcardially perfused for 2 min with PBS followed by 4% paraformaldehyde for 10 min. Brains were further fixed in 4% paraformaldehyde overnight and equilibrated successively in 15 and 30% sucrose in PBS for cryoprotection. Tissues were then frozen in dry ice and embedded in Tissue Plus optimum cutting temperature medium (Fisher HealthCare, Houston, TX). Cryostat sections of 30 μm were collected and placed into 24-well plates with antifreeze (30% ethyleneglycol, 30% glycerol, 10% 2 X PO_4_ buffer (0.244M) in dH_2_0). Prior to staining, free floating sections were washed 2 × 5 min in PBS and incubated for 1 hr at room temperature with blocking solution (1% bovine serum albumin (Gemini Bio Products, West Sacramento, CA) in PBS). Sections were then incubated overnight at 4°C with the following antibody dilutions in blocking solution: anti-Iba1 (1:1000; Thermo Fisher Scientific, Waltham, MA), anti-P2RY12 (1:1000; AnaSpec, Fremont, CA), anti-GFAP (1:500; Abcam, Cambridge, United Kingdom), anti-AOAH (1:100; Santa Cruz Biotechnology, Dallas, TX), and anti-IL-4 (1:500; Thermo Fisher Scientific). Brain sections were then washed 4 × 5 min with PBS followed by overnight incubation at 4°C with the following secondary antibody dilutions in blocking solution: Alexa 488-goat anti-rabbit (1:1000; Thermo Fisher Scientific) and Alexa 594-donkey anti-mouse (1:1000; Thermo Fisher Scientific). Secondary antibodies were washed off with PBS (2 × 5 min), incubated for 10 min in PBS/10 ug/mL 4’-6-diamidino-2-phenylindole (DAPI, Thermo Fisher Scientific) to stain nuclei, and further washed in PBS (3 × 5 min). Sections were then mounted onto gel-coated slides, dried for 30 min, and slides were mounted with 60 × 22 mm cover glass (VWR, Radnor, PA) using Clear-Mount Mounting Solution (Thermo Fisher Scientific) prior to imaging.

### PLX5622 treatment

For microglial elimination, colony-stimulating factor-1 receptor (CSF1R) inhibitor PLX5622 was purchased from MedChem Express (Monmouth Junction, NJ). The drug was dissolved in 10% DMSO and 90% corn oil, and AOAH-deficient mice were gavaged with 90 mg/kg of drug solution for 5 d. WT mice were gavaged with 10% DMSO and 90% corn oil only as a control. After 5 d of treatment, we ceased drug administration and allowed microglia to repopulate the CNS.

### Pelvic allodynia

Pelvic allodynia in WT, *Aoah*−/−, and *Aoah*−/−/*Tlr4*−/− mice was measured by quantifying von Frey filament stimulation to the pelvic region, as previously described in Rudick et al., 2012 [45]. Briefly, untreated and treated mice were singly placed in a test chamber and allowed to acclimate to their environment for 5 min. Five von Frey filaments (lowest-highest force) were applied 10 times to the pelvic region and responses were recorded. An animal was considered to be in pain if it jumped, lifted and shook the hind paws, or excessively licked the pelvic region.

### Cytokine array

The Proteome Profiler Mouse Cytokine Array Panel A (R&D Systems, Minneapolis, MN) was used to analyze cytokine expression in WT and *Aoah*−/− cortex following the protocol provided by the manufacturer. WT and *Aoah*−/− cortical lysates were prepared by homogenizing dissected cortical tissue in buffer consisting of 50 mM Tris HCl (pH 8.0), 150 mM NaCl, 1 mM EDTA, 0.5% sodium deoxycholate, and protease inhibitor cocktail (Millipore Sigma). After homogenization, 1% NP-40, 0.1% SDS, and 1% Triton were added to each sample and incubated on ice for 2 hrs followed by centrifugation and protein determination. Densitometry was done using ImageJ 1.52q software (National Institute of Health and the Laboratory for Optical and Computational Instrumentation).

### Western blotting

The immortalized murine microglial cell line, BV2 (Elabscience, Wuhan, China), were utilized for Western blotting. BV2 cells were plated at 500,000 cells/well for 48 hrs. Cells were then activated with lipopolysaccharides (LPS) from *Escherichia coli* 055:B5 (1μg/mL; Millipore Sigma, Burlington, MA) or with heat-killed stool slurry from WT or AOAH-deficient mice homogenized in PBS (1 mg/mL) for 0, 0.5, 1, 2, 6, or 24 hrs. Cells were then lysed for 30 min using RIPA buffer consisting of 50 mM Tris HCl (pH 8.0), 150 mM NaCl, 1 mM EDTA, 1% NP-40, 0.5% sodium deoxycholate, 0.1% SDS, 1% Triton, and protease inhibitor cocktail (Millipore Sigma). After centrifugation and protein determination, cell lysates or cell media were subjected to SDS-PAGE using 4-20% gradient Tris-glycine gels (Bio-Rad; Hercules, CA) followed by transfer to Immobilon P membranes (EMD Millipore). Following transfer, the membranes were incubated for 30 min in blocking buffer consisting of 10 mM Tris-HCl (pH 8.0), 150 mM NaCl, 0.01% Tween-20 (TBST), and 3% non-fat dry milk (Cell Signaling, Danvers, MA). Membranes were then incubated overnight at 4°C with the following primary antibodies diluted in blocking buffer: anti-CD11b (1:2000; Novus Biologicals, Littleton, CO), anti-TNFα (1:2000; Proteintech, Rosemont, IL), anti-CD68 (1:2000; Abcam), and anti-β-Actin (1:5000; Santa Cruz Biotechnology). After incubation, membranes were washed 5 × 5 min with TBST, incubated in blocking buffer for 30 min, and then incubated for 90 min with either horse-radish peroxidase (HRP) conjugated goat anti-rabbit IgG (1:10000; EMD Millipore) or HRP conjugated goat anti-mouse IgG (1:10000; Thermo Fisher Scientific). Blots were washed 5 × 5 min with TBST, and processed for chemiluminescence using SuperSignal West Dura Extended Duration Substrate (Thermo Fisher Scientific) prior to developing. All densitometric analyses were done using ImageJ 1.52q software (National Institute of Health and the Laboratory for Optical and Computational Instrumentation).

### Skeletal analyses

Morphological analysis of microglia was quantified by skeletal analyses and closely followed the protocol previously published by Young & Morrison [62]. Briefly, coronal brain sections (30 μm) from WT and AOAH-deficient mice were stained for the microglial marker P2RY12 and DAPI to label nuclei (as described in the section **Immunohistochemistry**). Photomicrographs taken with a 20X objective were processed to a binary image and skeletonized using Image J. Skeletonized images were then run through the AnalyzeSkeleton (2D/3D) plugin to quantify microglial morphology. Outliers that were considered noise were removed prior to data analyses.

### Transepithelial electrical resistance

Cecum permeability was measured as transepithelial electrical resistance (TEER) as previously described in Rahman-Enyart et al. [42]. Briefly, ceca were bisected and placed into Ringer’s solution prior to mounting on cassettes with an aperture of 0.126 cm^2^. Cassettes were then placed inside an Ussing chamber (Physiologic Instruments EM-CSYS-2) and KCl saturated salt-bridge electrodes were inserted. The chamber was maintained at 37°C with Ringer’s solution and bubbled with carbogen (95% O_2_/5% CO_2_) until resistance was stable (~1 hr). The voltage was then clamped and current was passed every 3 s at 30 s intervals via a VCC MC2 multichannel voltage-current clamp amplifier run by Acquire and Analyze software v.2.3.4 (Physiologic Instruments).

### Statistical analyses

Results are presented as average ± SEM. Student’s t-test or one-way analysis of variance (ANOVA) followed by Tukey’s Multiple comparisons test were utilized for data analyses. All statistical tests were run through Prism software, version 6 (GraphPad, Inc). Results between-groups were considered statistically significant at P<0.05.

## RESULTS

### AOAH is expressed in cortical microglia

We have previously reported AOAH expression in NeuN-positive neurons as well as Purkinje cells of the cerebellum [61]. To identify whether AOAH is also expressed in glial cells, we performed immunohistochemistry on cortical sections from WT mice. We were able to observe cortical microglia by immunostaining for ionized calcium binding adaptor molecule 1 (Iba1) and purinergic receptor P2Y12 (P2RY12), with complete colocalization and more robust staining with anti-P2RY12 (Fig. 1A-C). Since we observed brighter cell bodies and processes when immunostaining for P2RY12 compared to Iba1 (Fig. 1B compared to 1A), we utilized anti-P2RY12 to identify microglia for the remainder of our experiments.

**Figure 1.**
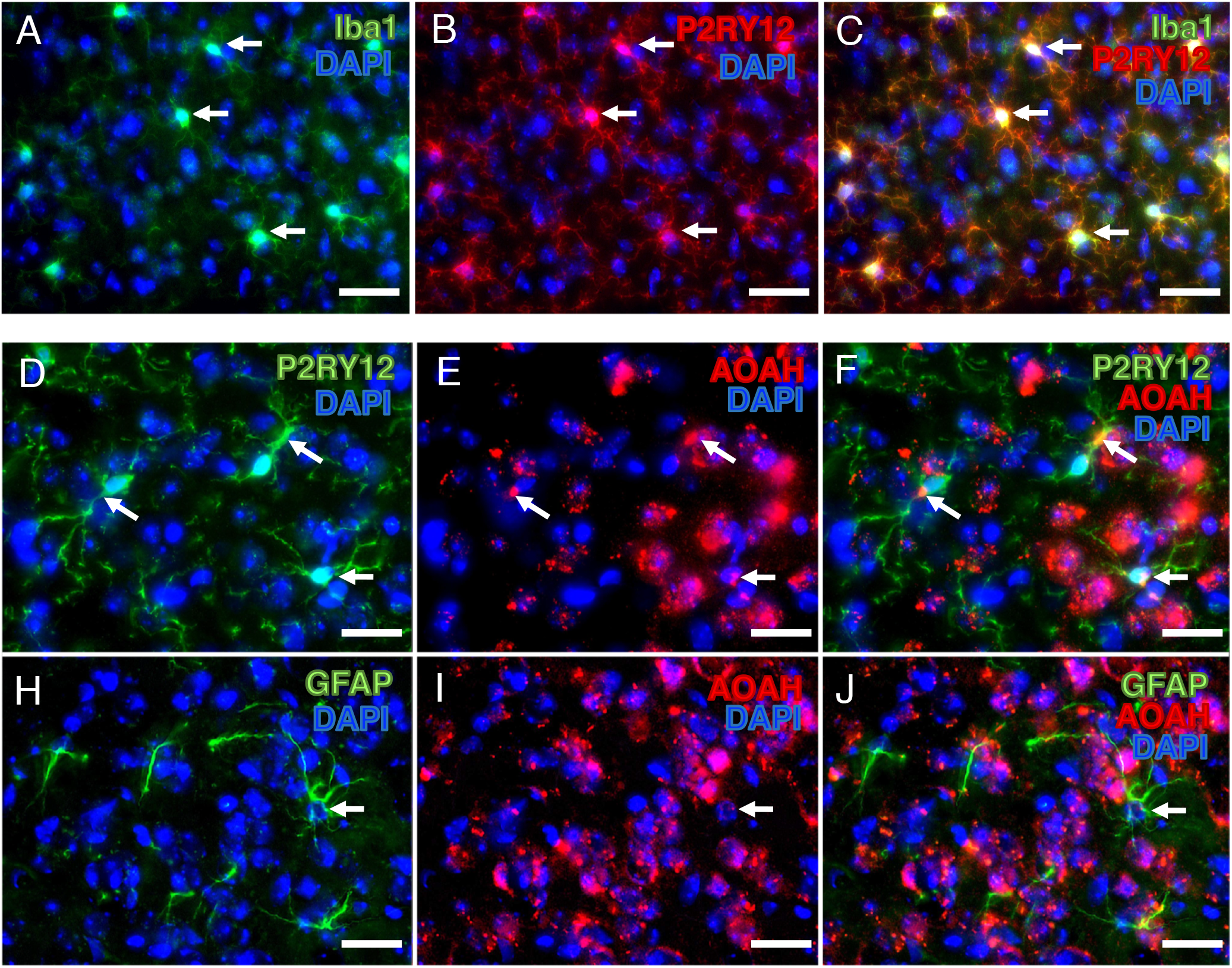
Immunostaining of AOAH in glial cells. **A-C:** WT mouse prefrontal cortex was stained for the microglial protein P2RY12 (green) and AOAH (red). DAPI staining nuclei shown in blue (scale bar: 30 μm). **D-F:** WT mouse prefrontal cortex was stained for the astrocyte protein glial fibrillary acidic protein (GFAP, green) and AOAH (red). DAPI staining nuclei shown in blue (scale bar: 30 μm).

Immunohistochemistry in cortical sections revealed AOAH expression in P2RY12-positive microglia, mainly within the perinuclear region (Fig. 1D-F). In contrast, glial fibrillary acidic protein (GFAP)-positive astrocytes showed low to background expression of AOAH protein (Fig. 1H-J). Our initial findings indicate that AOAH may play a biological role in microglia and possibly in microglial activation.

### Eliminating CNS microglia improves pelvic allodynia in AOAH-deficient mice

Microglial activation is a well-established player in the pathogenesis of chronic pain [16]. Our lab has identified that mice deficient for AOAH exhibit symptoms and comorbidities similarly observed in patients with IC/BPS, including pelvic pain [61]. To identify whether microglia may play a role in the pelvic pain phenotype observed by AOAH deficiency, we used a pharmacological approach to eliminate microglia from the CNS followed by analysis of pelvic allodynia. Microglia were ablated in AOAH-deficient mice via gavage of the CSF1R inhibitor PLX5622 (90 mg/kg) for 5 days. Allodynia was quantified in response to von Frey filaments applied to the pelvic region. Similar to our previous observations [42,61], we observed that baseline allodynia in AOAH-deficient mice was significantly elevated compared to WT (Fig. 2). AOAH-deficient mice that received PLX5622 treatment showed a 45% reduction in pelvic allodynia compared to baseline AOAH-deficient mice in response to the highest stimulus (Fig. 2, 88.13 ± 4.11% response at baseline vs 46.25 ± 9.30% response after treatment). A 34% reduction was observed after PLX5622 treatment in response to the second highest stimulus (Fig. 2, 71.88 ± 7.08% response at baseline vs 47.50 ± 8.59% response after treatment). After microglial depletion in AOAH-deficient mice, we withdrew treatment for 5 days to allow for repopulation of CNS microglia as previously described in Rice et al [44]. Microglial repopulation resulted in a significant increase in pelvic allodynia compared to treated animals in response to the three highest stimuli (71.82 ± 6.30%, 80.91 ± 6.67%, 88.18 ± 4.00% response from lowest to highest filament), similar to our observations at baseline for AOAH-deficient mice (Fig. 2). These data suggest that microglia, in part, play a role in the pelvic pain phenotype observed in AOAH-deficient mice.

**Figure 2.**
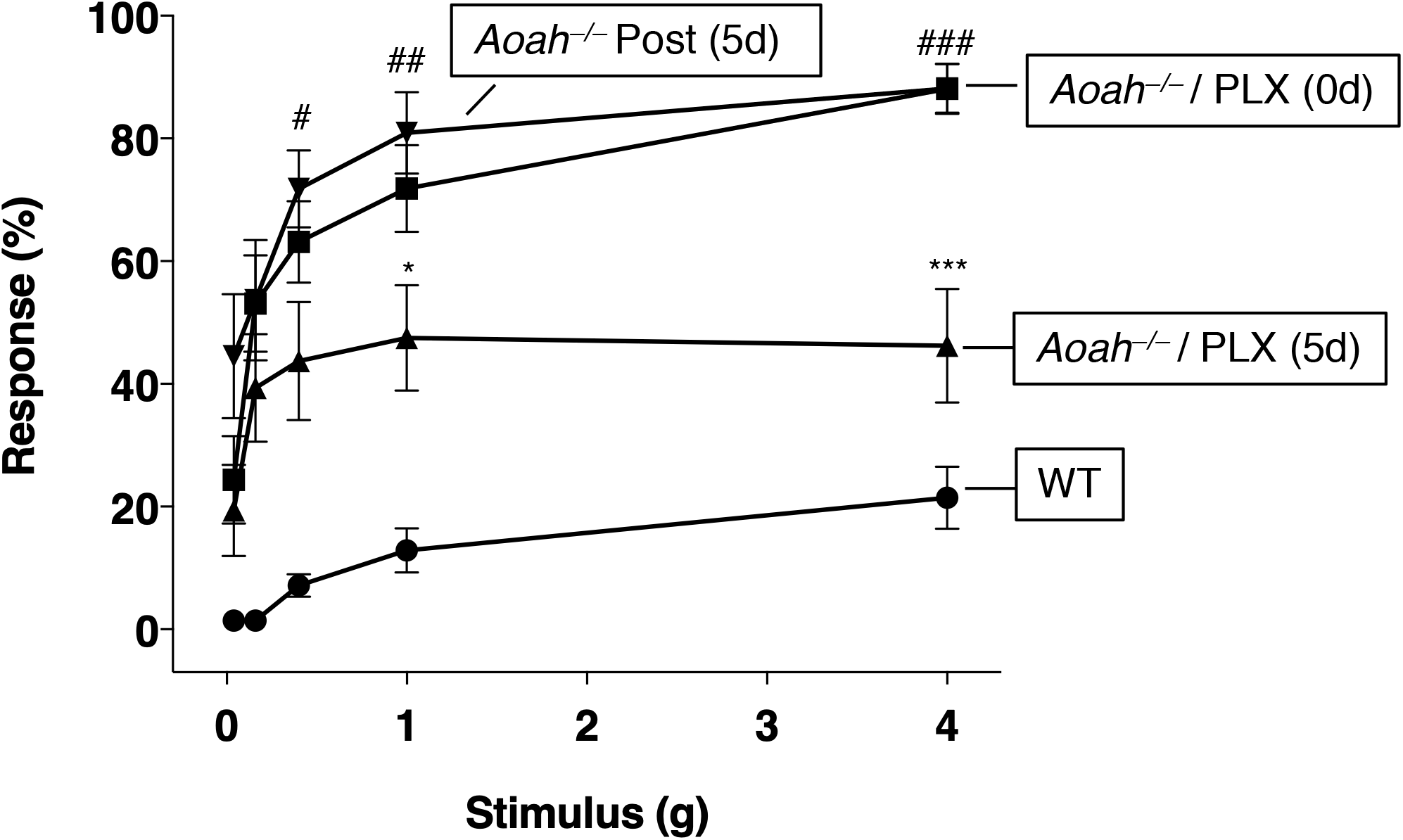
Pharmacologic ablation of microglia improves pelvic allodynia in AOAH-deficient mice. Stimulating the pelvic region with von Frey filaments revealed increased response in untreated and post-treated AOAH-deficient mice compared to WT and AOAH-deficient mice that were administered 90mg/kg of PLX5622 for 5 days by oral gavage in order to eliminate central nervous system (CNS) microglia (n=7 mice for WT, n=16 for baseline and PLX5622-treated AOAH-deficient mice, n=11 mice for AOAH-deficient mice post-PLX5622 treatment; *P<0.05, ***P<0.001 PLX5622 treatment compared to AOAH-deficient baseline, #P<0.05, ##P<0.01, ###P<0.001 Post (5 days) PLX5622 treatment compared to AOAH-deficient baseline, One-Way ANOVA followed by post-hoc Tukey HSD). Data represented as average response (%) ± SEM.

### Cytokine expression is upregulated in AOAH-deficient cortex

Activated microglia are a source of cytokine release which can drive immune responses and other neuromodulatory functions [23]. To address levels of brain cytokine expression in AOAH-deficient mice, we utilized a mouse cytokine array panel against 40 mouse cytokines. We observed several pro- and anti-inflammatory cytokines that were upregulated in AOAH-deficient cortex compared to WT, including IL-4 (Fig. 3). In addition, we also observed a down regulation of several cytokines in AOAH-deficient cortex, suggesting a complex but unique biochemical phenotype in AOAH-deficient cortex compared to WT.

**Figure 3.**
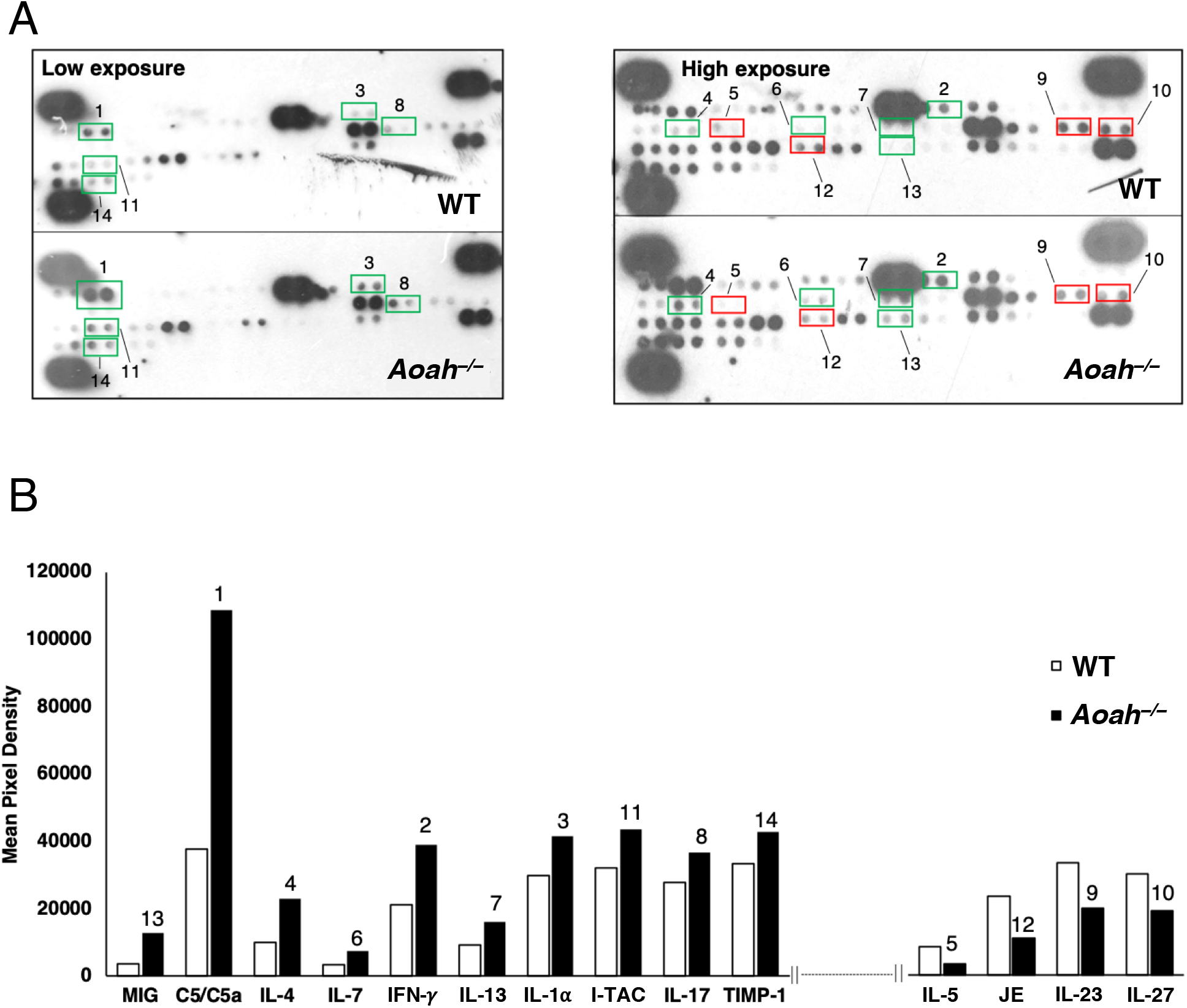
AOAH-deficient mice exhibit altered cytokine expression in the prefrontal cortex. **A:** Cytokine expression of WT and AOAH-deficient prefrontal cortex was analyzed using Proteome Profiler – Mouse Cytokine Array Panel A (R&D Systems, #ARY006). **B:** Cytokine expression data was quantified by densitometry (reported as mean pixel density).

To corroborate our array findings and identify whether aberrant cytokine expression in AOAH-deficient mice was observed in microglia, we utilized immunohistochemistry to identify colocalization between the anti-inflammatory cytokine IL-4 and the microglia specific protein P2RY12. We observed low levels of IL-4 expression in microglia in WT prefrontal cortex (Fig. 4, top row). AOAH deficiency resulted in increased microglia IL-4, which was observed in both the cell body and processes of microglial cells (Fig. 4, middle row).

**Figure 4.**
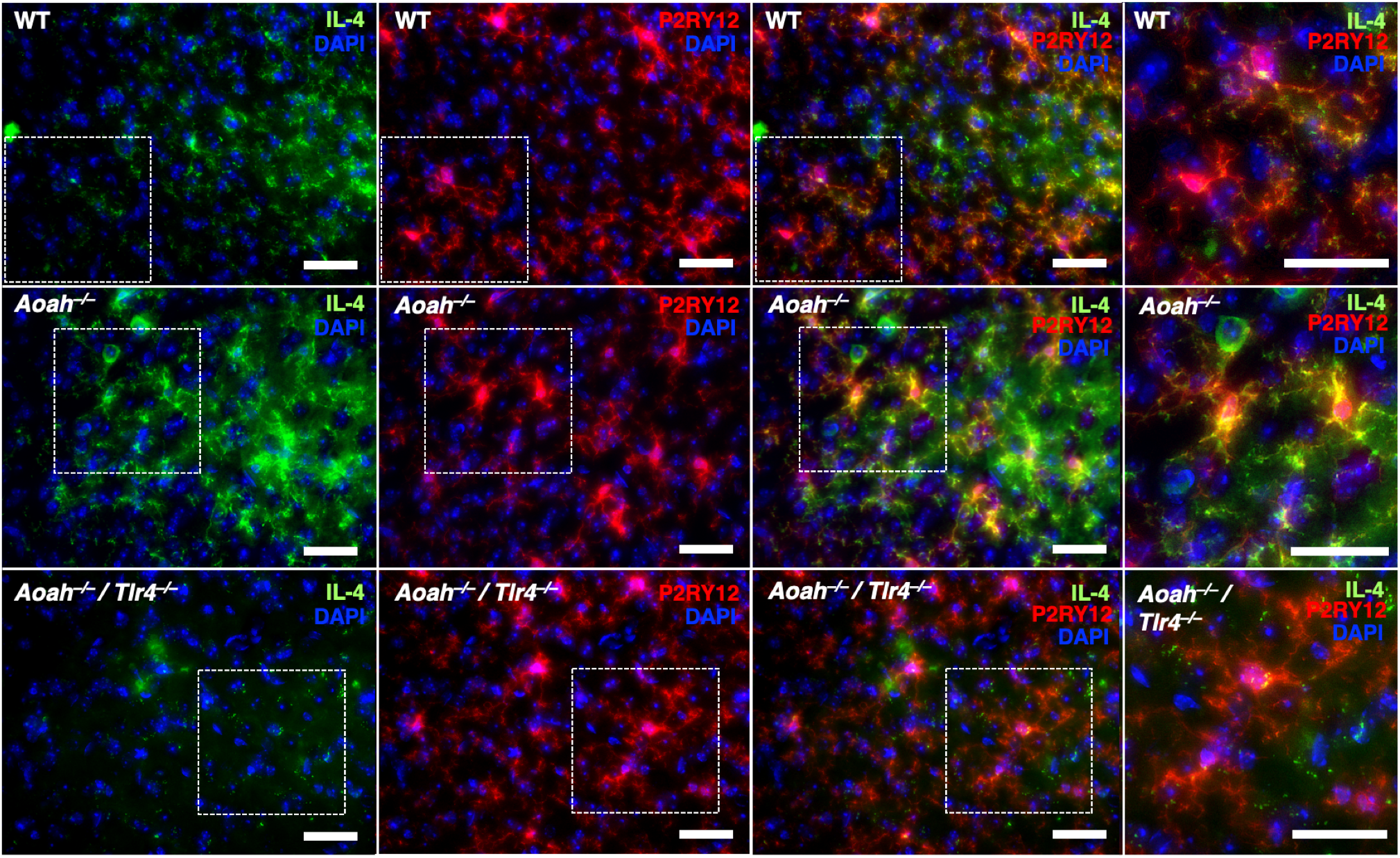
Immunostaining of IL-4 in cortical microglial cells. Mouse prefrontal cortex from WT (top row), *Aoah*−/− (middle row), and *Aoah*/*Tlr4*−/− (bottom row) mice were stained for the anti-inflammatory cytokine IL-4 (green) and AOAH (red). DAPI staining nuclei shown in blue (scale bar: 30 μm).

Toll-like receptors (TLRs), including TLR4, play an important role in immune response through the recognition of microbial components such as bacterial lipopolysaccharides (LPS) [27]. Since AOAH-deficient mice exhibit a “leaky gut” phenotype, circulating microbial products may be transported to the brain via the vagus nerve or a disrupted blood brain barrier where they have the ability to activate microglial TLR4 [16,42]. To observe whether TLR4 play a role in microglial activation in AOAH-deficient mice, we analyzed IL-4 expression in mice that were deficient for both AOAH and TLR4. Deficiency of both AOAH and TLR4 resulted in low levels of microglial IL-4 expression, similar to our observations in WT (Fig. 4, bottom row compared to top row). These data suggest that TLR4 activation plays a role in the microglial activation regulated by AOAH deficiency.

### Differential activation of BV2 cells by stool slurry from WT and AOAH-deficient mice

Since circulating microbes may be responsible for TLR4 activation in AOAH-deficient mice, we next sought to quantify differences in activation in a microglial cell line in response to gut bacteria. Activating BV2 cells with LPS from *E. coli* (1μg/mL) resulted in increased activation marker CD11b in cell lysates, peaking at 6 hrs of activation, and pro-inflammatory cytokine TNFα in cell media peaking at 1 hr of activation (Fig. 5B and C respectively). These data show that BV2 cells are activated by LPS in a time-dependent manner and show characteristics that mimic microglial activation.

**Figure 5.**
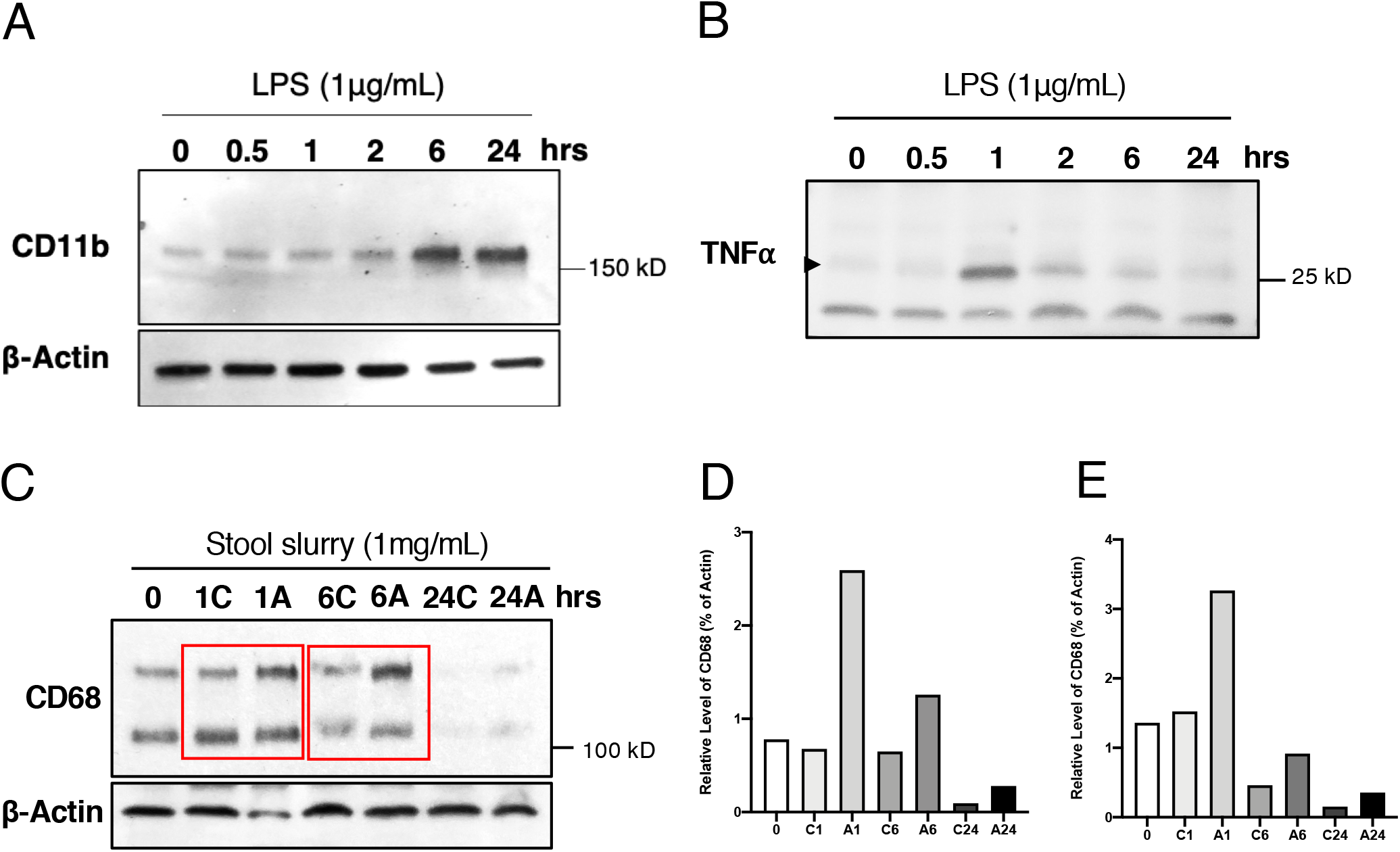
Microbiome-dependent activation of BV2 cells. **A:** Detergent extracts (50μg of protein/lane) from BV2 cells activated with 1μg/mL of LPS for 0, 0.5, 1, 2, 6, or 24 hours were analyzed by SDS-PAGE using 4-20% Tris-glycine gels followed by Western blotting. Blots were probed with CD11b (top panel, 1:2000) and β-Actin (bottom panel, 1:10000). **B:** Cultured media (50μg of protein/lane) from BV2 cells activated with 1μg/mL of LPS for 0, 0.5, 1, 2, 6, or 24 hours were analyzed by SDS-PAGE using 4-20% Tris-glycine gels followed by Western blotting. Blots were probed with TNFα (1:2000). **C:** Detergent extracts (50μg of protein/lane) from BV2 cells activated with 1mg/mL of stool slurry from WT (C) or AOAH-deficient (A) stool for 0, 1, 6, or 24 hours were analyzed by SDS-PAGE using 4-20% Tris-glycine gels followed by Western blotting. Blots were probed with CD68 (top panel, 1:2000) and β-Actin (bottom panel, 1:10000). **D and E:** Band intensities (from C) were quantified by densitometric analysis and reported as levels of the top (D) and bottom (E) bands of CD68 relative to levels of β-Actin.

We next determined whether BV2 cells responded differentially in response to bacteria from WT and AOAH-deficient gut by activating cells with stool slurry (1 mg/mL). Upon activation of BV2 cells with heat-killed stool slurry from WT mice, we observed cell activation peaking at 1 hr as quantified by activation marker CD68 expression (Fig. 5C-E). In comparison, when activating with slurry from AOAH-deficient mice, cells showed greater activation at 1 hr and activation was sustained with high levels of CD68 after 6 hrs of activation (Fig. 5C-E). Since CD68 glycosylation correlates with macrophage activation [11], we quantified both observed CD68 bands and observed higher levels of both forms after activation with AOAH-deficient stool slurry at all time points (Fig. 5D and E).

### AOAH-deficient mice exhibit activated microglial morphology

Our data so far have suggested activated microglia in AOAH-deficient mice through addressing their biochemical phenotype (Figs. 3 and 4). Microglia are highly dynamic cells that exhibit morphological changes upon activation, where ramified microglia are considered to be in resting/surveillance-mode and less ramified (bushy or ameboid) microglia are considered to be activated [30].

In order to quantify microglial morphology, brain sections were stained with the microglial marker P2RY12 followed by image processing and skeletal analyses using ImageJ (Fig. 6A for example of photomicrographs analyzed). Skeletal analyses of microglia in the prefrontal cortex revealed that AOAH-deficient mice exhibit microglia with fewer branches (Fig. 6B, 59.82 ± 3.45 for *Aoah*−/− vs 78.62 ± 4.98 for WT), fewer endpoints (Fig. 6C, 48.14 ± 1.73 for *Aoah*−/− vs 63.71 ± 3.17 for WT), and shorter processes (Fig. 6D and E, 42.19 ± 2.27 μm and 59.52 ± 2.65 μm for *Aoah*−/− vs 57.40 ± 4.42 μm and 82.11 ± 5.73 μm for WT, respectively) compared to WT cortical microglia. We did not observe any changes in the number of microglia between AOAH-deficient and WT cortical microglia (Fig. 6F, 41.80 ± 1.27 for *Aoah*−/− vs 38.27 ± 1.96 for WT). These data suggest that AOAH-deficient mice exhibit an activated microglial morphology in the prefrontal cortex.

**Figure 6.**
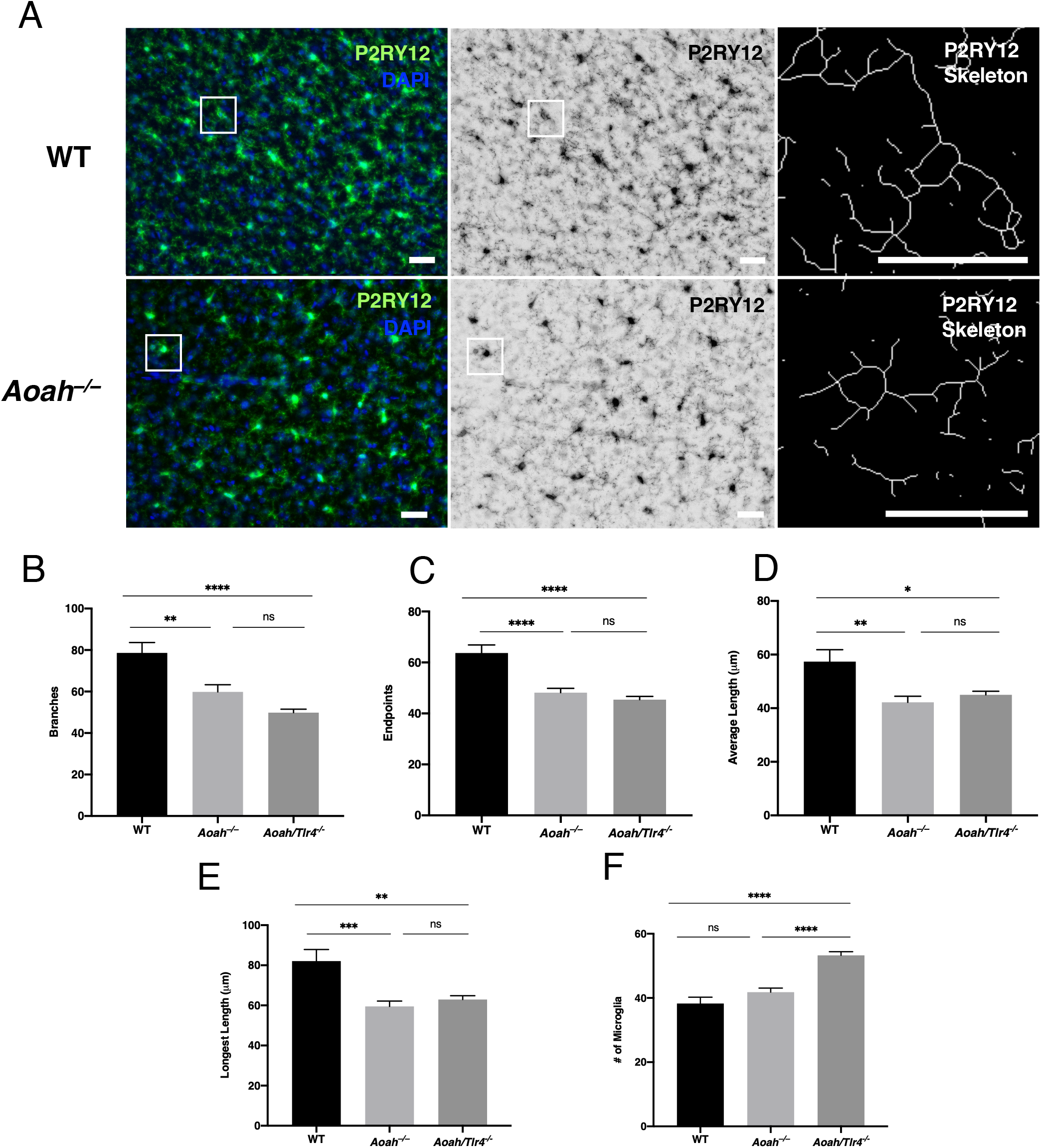
AOAH-deficient prefrontal cortex exhibits an activated microglia phenotype. **A:** Example of photomicrographs used for skeletal analyses. Left column shows immunostaining of P2RY12 (green) in cortical microglial cells in WT (top) and AOAH-deficient (bottom) mice. DAPI staining nuclei shown in blue (scale bar: 30 μm). Middle column shows 8-bit grayscale images (scale bar: 30 μm). Right column shows example of skeletonized microglia used for quantification (scale bar: 30 μm). **B-F:** Quantified microglia morphology of prefrontal cortex by skeletal analyses. *Aoah*−/−and *Aoah*/*Tlr4*−/− cortical microglia exhibit fewer branches (B), fewer endpoints (C), shorter length of their average process (D), and shorter length of their longest process (E), consistent with an activated microglia phenotype. *Aoah*/*Tlr4*−/− mice have an increased number of prefrontal cortical microglia compared to wild-type and AOAH-deficient mice (F, n=10 for all groups; *P<0.05, **P<0.01, ***P<0.001, ns = not significant, One-Way ANOVA followed by posthoc Tukey HSD). Data represented as average ± SEM.

We also sought to identify whether TLR4 played a role in the activated microglial phenotype observed in AOAH-deficient mice, since we observed decreased IL-4 expression in *Aoah/Tlr4*−/− cortical microglia (Fig. 4). In contrast to our findings showing decreased IL-4 expression, *Aoah/Tlr4*−/− mice exhibited morphology of activated microglia in the prefrontal cortex (Fig. 6B-F, 49.82 ± 1.68 for number of branches, 45.40 ± 1.31 for number of endpoints, 44.99 ± 1.35 μm for length of processes, and 62.96 ± 1.86 μm for length of longest process) including an increase in the number of microglia (Fig. 6F, 53.27 ± 1.14). These findings suggest that AOAH deficiency is responsible for the activated microglial phenotype independent of TLR4.

Since AOAH-deficient mice exhibit pelvic pain [61] and we observed activated microglia in the prefrontal cortex (Figs. 4 and 6), a brain region associated with neuropathic pain, we next sought to analyze microglial activation in another brain region responsible for pain modulation, the paraventricular nucleus (PVN). Similar to our observations in the prefrontal cortex, AOAH-deficient microglia in the PVN exhibited altered microglial morphology as observed by decreased number of branches (Fig. 7A, 66.12 ± 3.98 for *Aoah*−/− vs 102.80 ± 5.81 for WT) and endpoints (Fig. 7B, 52.37 ± 3.07 for *Aoah*−/− vs 64.13 ± 3.60 for WT). We did not observe differences in the length of processes (Fig. 7C and D, 44.15 ± 3.87 μm and 64.10 ± 4.66 μm for *Aoah*−/− vs 43.27 ± 4.53 μm and 64.62 ± 6.48 μm for WT, respectively) or number of microglia in PVN microglia between WT and AOAH-deficient mice (Fig. 7E, 32.71 ± 3.23 for *Aoah*−/− and 28.25 ± 1.72 for WT). Overall, these data show activated microglia morphology in AOAH-deficient mice in brain regions associated with neuropathic pain.

**Figure 7.**
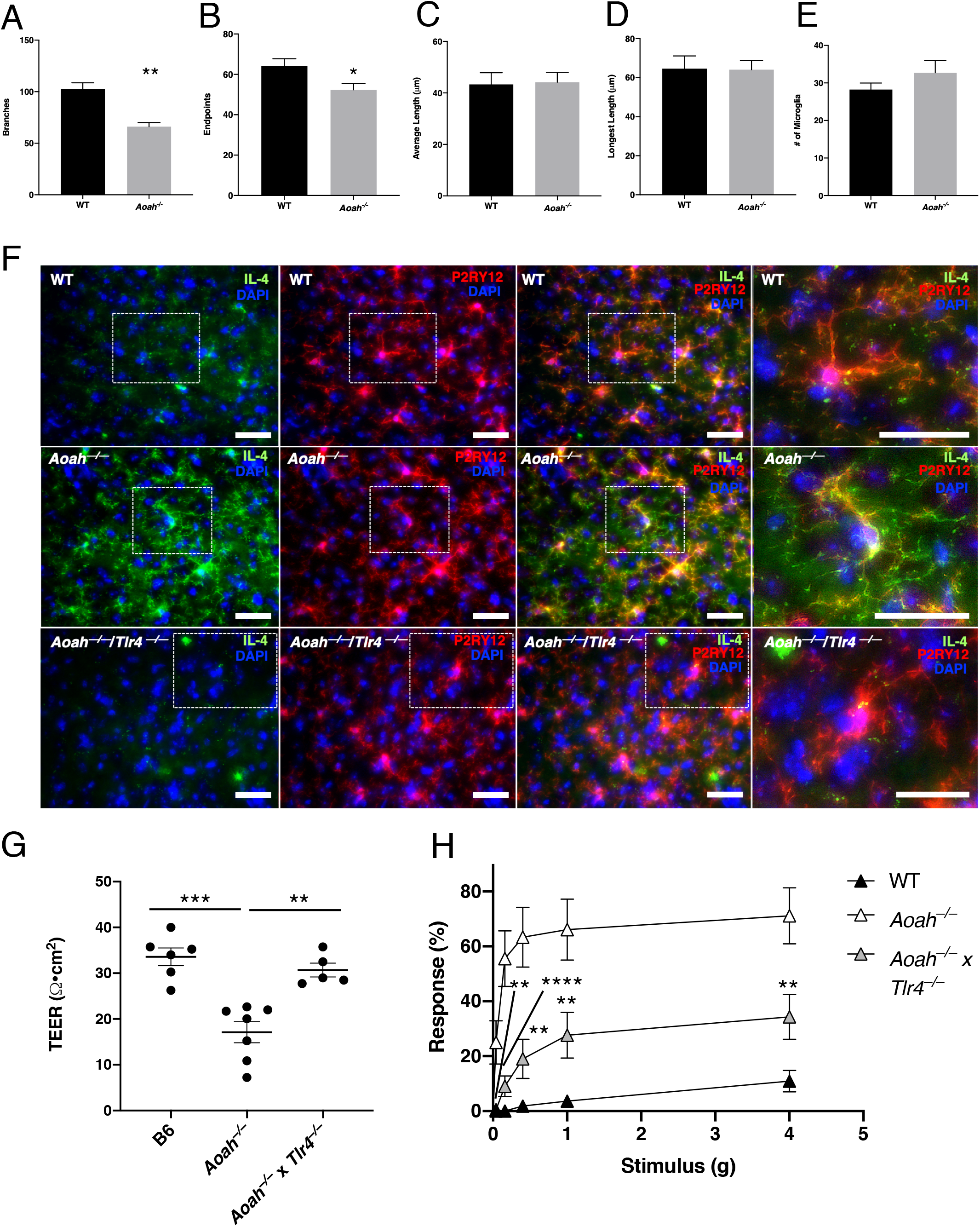
AOAH-deficient paraventricular nucleus (PVN) activated microglia and TLR4 regulates AOAH-deficient microglial activation and pelvic pain. **A-E:** Quantified microglia morphology of PVN by skeletal analyses. *Aoah*−/− PVN microglia exhibit fewer branches (A) and fewer endpoints (B) compared to WT, consistent with an activated microglia phenotype (n=5 for WT, n=6 for *Aoah*−/−; *P<0.05, **P<0.01, Student’s t test, two tailed). The average length of processes (C), length of longest process (D), and number of microglia (E) was not significantly different between groups. Data represented as average ± SEM. **F:** Immunostaining of IL-4 in PVN microglial cells. Mouse PVN from WT (top row), *Aoah*−/− (middle row), and *Aoah*/*Tlr4*−/− (bottom row) mice were stained for the anti-inflammatory cytokine IL-4 (green) and AOAH (red). DAPI staining nuclei shown in blue (scale bar: 30 μm). **G:** Ceca of female *Aoah*−/− mice showed a decrease in transepithelial electrical resistance (TEER) compared to WT and *Aoah*/*Tlr4*−/− mice (WT mice: n=6, *Aoah*−/− mice: n=7, *Aoah*/*Tlr4*−/−: n=5 **P<0.01, ***P<0.001, One-Way ANOVA followed by post-hoc Tukey HSD). Data represented as individual values and average (represented by horizontal line) ± SEM. **H:** AOAH-deficient mice exhibited increased pelvic allodynia compared to WT, which was significantly alleviated in *Aoah*/*Tlr4*−/− mice as shown through response to von Frey filaments stimulating the pelvic region (WT mice: n=11, *Aoah*−/− mice: n=18, *Aoah*/*Tlr4*−/−: n=21; **P<0.01, ****P<0.0001, One-Way ANOVA followed by post-hoc Tukey HSD). Data represented as average response (%) ± SEM.

We next sought to analyze whether the PVN exhibited increased microglial IL-4 in the PVN, similar to our observations in the prefrontal cortex (Fig. 4). Through immunohistochemistry, we identified colocalization between IL-4 and P2RY12-positive microglia. Similar to our findings in the prefrontal cortex we observed low levels of IL-4 in WT PVN and high levels of IL-4 in AOAH-deficient PVN microglia, consistent with an activated microglia phenotype (Fig. 7F, top row compared to middle row). In addition, mice that were deficient for both AOAH and TLR4 showed low levels of IL-4 in the PVN, similar to our observations in WT, suggesting a role for TLR4 in microglial activation in AOAH-deficient mice (Fig. 7F, bottom row compared to top row).

### TLR4 regulates gut leakiness and pelvic allodynia in AOAH-deficient mice

Our data has shown that TLR4 modulates microglial activation, but not microglial morphology, in AOAH-deficient mice (Figs. 4, 6, and 7). Since we observed improved pelvic allodynia upon microglial elimination (Fig. 2), we next sought to identify whether TLR4 deficiency would improve symptoms observed in AOAH-deficient mice.

We have previously shown compromised gut epithelia in AOAH-deficient mice [42]. Consistent with these previous findings, we measured transepithelial electrical resistance (TEER) of ceca and observed reduced TEER in AOAH-deficient mice (Fig. 7G), suggesting a “leaky gut” phenotype. In contrast, ceca from mice that were deficient for both AOAH and TLR4 exhibited TEER comparable to WT mice, suggesting that TLR4 deficiency improves the “leaky gut” phenotype observed in AOAH-deficient mice (Fig. 7G).

AOAH-deficient mice exhibit increased pelvic allodynia (Fig. 2 and [61]). To examine whether TLR4 plays a role in the pelvic pain phenotype observed in AOAH-deficient mice, we measured pelvic allodynia in *Aoah/Tlr4*−/− mice by quantifying responses to von Frey filaments applied to the pelvic region. Mice that were deficient for both AOAH and TLR4 exhibited reduced pelvic allodynia compared to AOAH-deficient mice at all filaments examined (average 18.10 ± 5.27% total response for *Aoah/Tlr4*−/− mice vs 56.22 ± 9.31% total response for *Aoah*−/− mice), suggesting that TLR4 plays a role in the pelvic pain phenotype due to AOAH deficiency (Fig. 7H). A 67% reduction in total pelvic allodynia was observed in *Aoah/Tlr4*−/− compared to *Aoah*−/− mice. Deficiency of TLR4 in AOAH-deficient mice, however, did not completely alleviate the pelvic pain phenotype to levels observed in WT (Fig. 7H, average 3.46 ± 1.38% total response for WT). These data suggest that TLR4, in part, plays a role in pelvic allodynia of AOAH-deficient mice.

## DISCUSSION

We previously reported that AOAH-deficiency mimics several aspects of IC/BPS, including pelvic pain and gut dysbiosis [1,2,42,61]. Here we identified that activated microglia play a role in the pelvic pain phenotype observed in AOAH-deficient mice. Prior studies have implicated both microglial activation and dysbiosis of gut flora as regulators of chronic pain [5,16,20,22,26]. We observed that AOAH-deficient mice exhibited activated microglia in the prefrontal cortex and PVN, as observed by increased cytokine expression, such as IL-4, and microglial morphology [Figs. 4, 6, and 7]. Microglial release of IL-4, but not microglial morphology, relied on the expression of TLR4 [Figs. 4, 6, and 7]. These findings are consistent with a role for AOAH deficiency in “priming” microglia to a state allowing activation and TLR4 in microglial cytokine release and response (Fig. 8).

**Figure 8.**
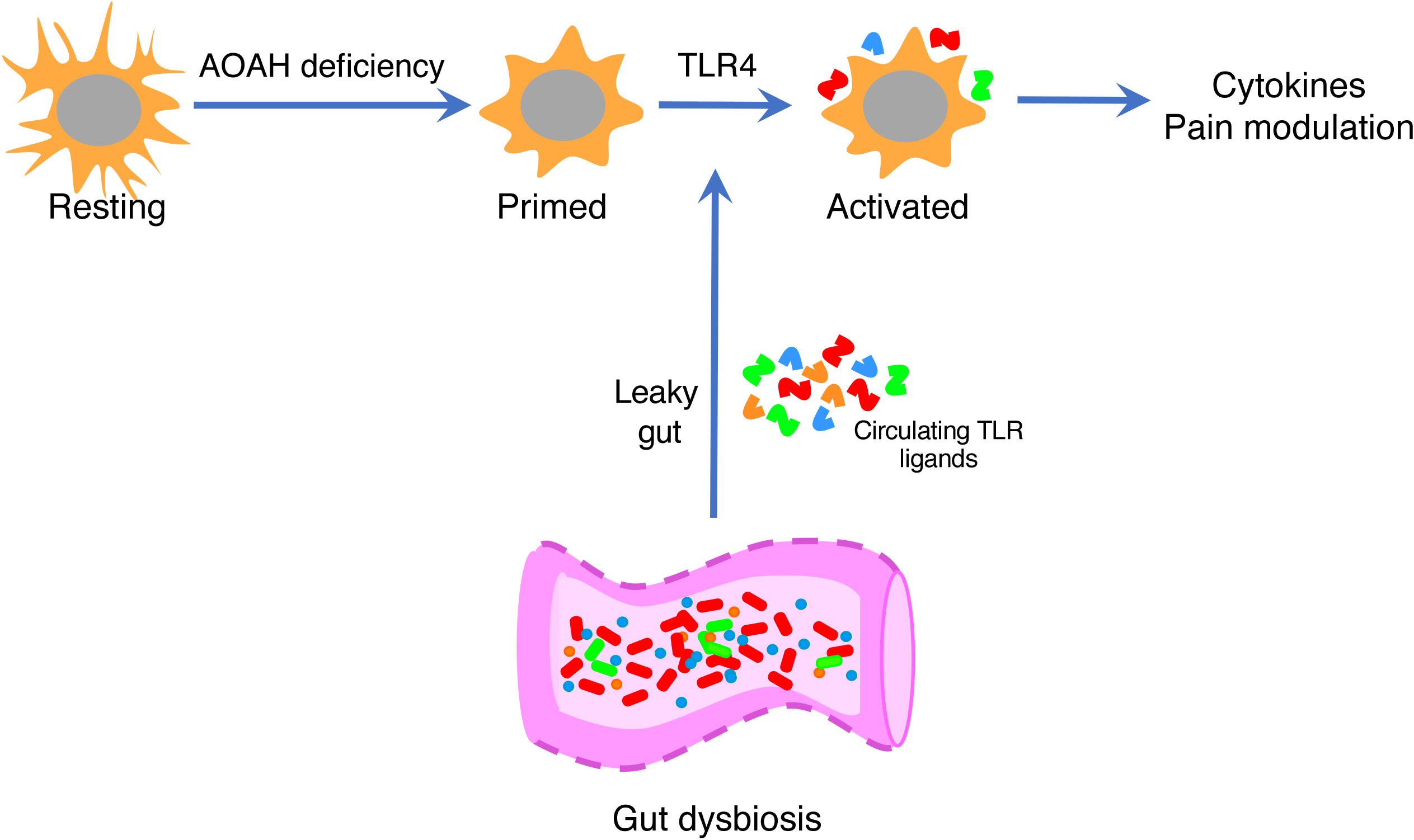
Model of microglial activation regulated by AOAH deficiency and TLR4. At baseline, microglia exhibit a ramified/resting phenotype. Deficiency of AOAH results in microglia with a primed phenotype. AOAH-deficient microglia can then become further activated (altered cytokine expression) by CNS TLR4. One possible mechanism of TLR4 activation could be by circulating microbial ligands from the gut crossing the blood brain barrier and activating primed microglia. In addition, the gut microbiome can alter vagus nerve signaling to the CNS. Prolonged microglial activation can then regulate pain modulation.

Our findings suggest that activated microglia in AOAH-deficient mice in-part modulates the pelvic pain phenotype, as pharmacological elimination of microglia improves pelvic allodynia in these mice (Fig. 2). As the gut microbiome can manipulate microglial function [18,36,55], we propose a functional model of microglial activation by gut components in AOAH deficiency-mediated pelvic pain (Fig. 8). AOAH-deficient mice exhibit gut dysbiosis and a compromised gut epithelia [42], allowing gut components including TLR ligands to enter the bloodstream. Circulating TLR ligands, such as bacterial LPS, can cross a compromised blood brain barrier (BBB) and directly activating primed microglia via TLR4 activation. Alternatively, altered microbiota can directly modify CNS physiology via the vagus nerve. In response to these inputs, altered microglial cytokine expression can then modify neural mechanisms of pain modulation (Fig. 8). One limitation of our proposed model is whether TLR ligands from the gut are responsible for AOAH-deficient microglial activation and, if so, whether these ligands cross to the brain via the vagus nerve or a compromised BBB. Further exploration between gut microbe-microglia association in AOAH-deficient mice is required to address these gaps.

TLR4 activation is a key player in regulating persistent pain, and blocking TLR functions have been advantageous for chronic inflammation and nerve injury models [7,9,10,41,60]. Previous studies have suggested a role for TLR4-mediated inflammatory response in IC/BPS, where peripheral blood mononuclear cells from IC/BPS patients showed higher levels of pro-inflammatory cytokines upon TLR stimulation, which correlated with pelvic pain severity [47–49]. We found that AOAH-deficient mice that were also deficient for TLR4 showed decreased microglial activation in the prefrontal cortex and PVN, improved cecum epithelial barrier function, and reduced pelvic allodynia compared to AOAH-deficient mice (Figs. 4 and 7). These findings align with those reported by Cui et al showing TLR4 dependency in bladder nociceptive response, spinal pro-inflammatory cytokine release, and spinal glial activation in an IC/BPS-like cystitis model [14]. Although the role of TLR4 in pelvic pain severity is compelling, whether microglial TLR4 is the key mediator is not well understood and will require future conditional deletions of *Tlr4* in microglia.

We observed altered expression of both pro- and anti-inflammatory cytokines in AOAH-deficient mice (Fig. 3). Pro-inflammatory cytokines released from glial cells or macrophages, such as TNFα, IL-1β, and IL-6 are well-studied regulators of disease severity observed in “classically activated” cells, whereas anti-inflammatory cytokines, such as IL-4, are considered to be tissue healing and present in “alternatively activated” cells [17,58,59,64]. One explanation for AOAH deficiency resulting in altered expression of both pro- and anti-inflammatory cytokines in the cortex may be due to a compensatory mechanism. A previous study demonstrated that IL-4 priming of mouse bone marrow-derived macrophages could induce a strong pro-inflammatory response upon activation of TLR4 via LPS [17]. Therefore, IL-4 expression not only implicates a “tissue protective” response, but also suggests a mechanism for increased pro-inflammatory cytokine release in response to challenges from leaked gut microbes and subsequent TLR4 activation in AOAH-deficient mice.

AOAH-deficient mice exhibit gut dysbiosis, including an enrichment in gut bacteria [42]. We have previously reported that microbiota manipulation can result in the alleviation of pelvic allodynia and anxious behavior in AOAH-deficient mice [42]. In addition, here we observed that exposure to WT or AOAH-deficient stool slurry resulted in differential activation of microglial BV2 cells *in vitro* (Fig. 5) and that microglial elimination improved pelvic pain (Fig. 2). These findings suggest a role for gut microbe-microglia interactions in pelvic allodynia, which may be regulated by gut composition. The gut microbiome of AOAH-deficient mice exhibits significantly increased abundance of several bacterial phyla compared to wild-type mice, including the presence of cyanobacteria [42]. A previous study by Mayer and colleagues exposed rat microglia *in vitro* with cyanobacterium *Oscillatoria* sp. LPS and observed a concentration-dependent release of both pro- and anti-inflammatory cytokines and chemokines [37]. In addition to increased cyanobacteria in AOAH-deficient microbiota [42], we also observed an upregulation of several pro- and anti-inflammatory cytokines in AOAH-deficient prefrontal cortex (Fig. 3). Therefore, the interaction between strains of cyanobacteria and microglia in AOAH-deficient mice may have an association with the observed pelvic pain phenotype.

We have previously shown *Aoah* as a genetic regulator of corticotropin releasing factor (CRF) in the PVN [2], a pertinent brain region for CRF-dependent pain modulation and stress response [31,54]. AOAH-deficient mice exhibit increased *Crf* expression as well as enhanced stress behaviors [2]. Microglia express receptors for CRF, and activation of microglia by CRF results in cell proliferation and cytokine release [29,57]. In addition, neuronal CRF release and microglial activation regulated by TLR4 signaling in the PVN have been observed in a rodent visceral hypersensitivity model of irritable bowel syndrome [63]. Since we have identified elevated *Crf* [2] and activated microglia in AOAH-deficient PVN (Fig. 7), it is possible that CRF-releasing neurons play a role in the primed microglial phenotype. The combination of leaked gut microbes (Fig. 8) and neuronal CRF may contribute to the persistent activation and diverse cytokine release of AOAH-deficient microglia.

Microglial activation has been observed in various clinical disorders including incidence of anxiety and depression [15,43,50], disorders that are comorbid with IC/BPS [12,24,38]. In addition, AOAH-deficient mice exhibit an anxiety/depressive phenotype [2]. Human studies have suggested microglial activation may play a role in depression. For example, postmortem brain analyses in depressed patients and suicide victims reveal increased microgliosis and a higher ratio of primed over ramified (resting) microglia [52,53,56]. Microgliosis has been observed in brains of suicide victims in several regions, such as the dorsolateral prefrontal cortex, anterior cingulate cortex, and mediodorsal thalamus [52,53]. In conjunction, positron emission tomography (PET) imaging in living human patients with depression have detected microglial activation correlating with suicidal ideation [25]. Whether microglial activation is associated with IC/BPS is currently unknown. Here we have shown microglial activation in a validated mouse model of IC/BPS, suggesting a similar phenotype may be observed in IC/BPS and similar mechanisms may mediate microglial pain modulation associated with pelvic pain in IC/BPS.

In summary, the data presented here show that AOAH-deficient mice exhibit activated microglia in pertinent brain regions for pain modulation, which is dependent on TLR4. These findings demonstrate TLR4 or other microglial regulators as promising pharmacological targets for treating pelvic pain disorders, such as IC/BPS.

## ETHICS APPROVAL AND CONSENT TO PARTICIPATE

All animals were maintained at the Center for Comparative Medicine at Northwestern University and utilized for experimentation under Northwestern IACUC approved protocols.

## COMPETING INTERESTS

The authors declare that they have no competing interests.

## FUNDING

This work was supported by NIH/NIDDK award R01 DK103769 (A.J.S., and D.J.K) and by NIH/NIDDK T32 DK062716 postdoctoral fellowship to A.R.-E.

## AUTHOR CONTRIBUTIONS

A.R.-E., A.J.S., J.L.B., D.R.W., and D.J.K. conceived and designed research; A.R.-E., R.E.Y., and W.Y. performed experiments and analyzed data; A.R.-E., R.E.Y., A.J.S., and D.J.K. interpreted results of experiments; A.R.-E. and D.J.K. prepared figures; A.R.-E. and D.J.K. drafted manuscript; A.R.-E., A.J.S., D.R.W., and D.J.K. edited and revised manuscript; A.R.-E., R.E.Y., W.Y., J.L.B., D.R.W., A.J.S., and D.J.K. approved final version of manuscript.

## ACKNOWLEDGEMENTS

We thank Dr. Robert Munford for generously providing AOAH-deficient mice and for many helpful discussions.

## Notes

### Competing Interest Statement

The authors have declared no competing interest.

